# Mitochondrial Dysfunction alters Early Endosome Distribution and Cargo Trafficking via ROS-Mediated Microtubule Reorganization

**DOI:** 10.1101/2024.08.21.608999

**Authors:** Anjali Vishwakarma, Lilia Chihki, Kiran Todkar, Mathieu Ouellet, Marc Germain

## Abstract

Mitochondria are essential for bioenergetic functions and various cellular processes, including differentiation and immunity, their dysfunction leading to several pathologies. While these pathologies have traditionally been associated with ATP deficits, mitochondrial dysfunction also leads to reactive oxygen species (ROS) generation, inflammation, and alterations the function of other organelles. While the negative impact of mitochondrial dysfunction on lysosomal activity has been established, the relationship between mitochondria and the rest of the endocytic compartment remains poorly understood. Here, we show that inhibiting mitochondrial activity through genetic and chemical approaches causes early endosome (EE) perinuclear aggregation and impairs cargo delivery to lysosomes. This impairment is due to altered microtubule architecture and centrosome dynamics, mediated by ROS. Antioxidants can rescue these EE defects, underlying the pivotal role of mitochondria in maintaining cellular activities through ROS regulation of microtubule networks. Our findings highlight the significance of mitochondria beyond ATP production, emphasizing their critical involvement in endocytic trafficking and cellular homeostasis. These insights emphasize mitochondria’s critical involvement in cellular activities and suggest novel targets for therapies to mitigate the effects of mitochondrial dysfunction.

## Introduction

Mitochondria are key bioenergetic organelles that play crucial roles in not only metabolite generation and exchange, but also in a range of cellular processes including cell differentiation and immunity (San-Millan, 2023). As a consequence, alterations in mitochondrial functions cause a wide range of muscular and neurological pathologies (Chen *et al*, 2022; Chen *et al*, 2023). While these pathologies were historically thought to result from ATP deficits, it is now clear that other mechanisms are also involved, including enhanced inflammation and the generation of reactive oxygen species (ROS) (Johnson *et al*, 2021; Nissanka & Moraes, 2018). In fact, ATP-independent roles of mitochondria are crucial for the control of cellular differentiation during development, inflammation and cancer (Klein *et al*, 2020).

At the cellular level, altered mitochondrial structure and function are linked to defects in other organelles. For example, loss of mitochondrial function leads to defects in lysosomal activity (Demers-Lamarche *et al*, 2016; Glancy *et al*, 2020). As lysosomes are the main degradative organelles, this affects protein turnover and causes the accumulation of aggregated material within the affected cells (Cox & Cachon-Gonzalez, 2012). Lysosomes receive their material to degrade through autophagy, which is known to be affected by mitochondrial dysfunction, and following endocytosis of extracellular material (de la Mata *et al*, 2016; Demers-Lamarche *et al*., 2016). In the endocytic pathway, extracellular material first accumulates in early endosomes (EEs) that sorts this material towards recycling to the cell surface or targets them to late endosomes and lysosomes for degradation (Naslavsky & Caplan, 2018). The sorting and transfer of material to recycling and late endosomes requires the action of small GTPases of the Rab family. Rabs coordinate all the steps required for cargo sorting and transfer to other endocytic organelles, including the transport along microtubules that allow fusion between vesicles and transfer to lysosomes (Homma *et al*, 2021).

While alterations in mitochondrial activity impact lysosomes, the consequences on other components of the endocytic pathway remain poorly understood. Here, we show that mitochondrial dysfunction disrupts EE distribution and cargo trafficking, revealing a previously unrecognized link between mitochondrial function and endocytic transport mechanisms. Specifically, the presence of genetic mutations in mitochondrial proteins or the chemical inhibition of the electron transport chain results in the perinuclear aggregation of EEs and impairs their ability to deliver cargo to lysosomes. These alterations in EE distribution are caused by changes in microtubule architecture and centrosome dynamics, which then facilitates EE transport and aggregation in the perinuclear region of the cell. We found that this is mediated by ROS, which affect microtubules and centrosome organization and thus EE distribution and function. Consequently, antioxidants rescue EE defects present in cells with mitochondrial dysfunction. Overall, our results demonstrate that ROS generated by defective mitochondria actively disrupt endocytic trafficking through the reorganisation of microtubule networks, underscoring the critical role of mitochondria in the maintenance of cellular activities.

## Results

### Mitochondrial dysfunction causes early endosome aggregation

Mitochondrial dysfunction impairs lysosomal activity, which is accompanied by the presence of enlarged and vacuolated lysosomes and late endosomes (Baixauli *et al*, 2015; Demers-Lamarche *et al*., 2016; Fernandez-Mosquera *et al*, 2019). We thus determined whether this extended to the rest of the endocytic compartment. For this, we first used mouse embryonic fibroblasts (MEFs) in which the essential mitochondrial protein OPA1 is deleted as a model for mitochondrial dysfunction (Demers-Lamarche *et al*., 2016; Patten *et al*, 2014). As we previously reported (Demers-Lamarche *et al*., 2016), these cells contained large vacuolated structures marked with the lysosomal marker LAMP1 (Figure 1A-B). As early endosomes (EEs) are required to deliver extracellular material to lysosomes, we determined whether they become vacuolated similar to lysosomes by staining cells with an antibody against the EE marker Rab5. However, in contrast to LAMP1-positive lysosomes, Rab5-positive EEs did not appear vacuolated in OPA1 KO MEFs (Figure 1A-B). Nevertheless, there was a clear shift in EE distribution in OPA1 KO cells. Whereas EEs were equally dispersed throughout in control cells, we observed perinuclear clustering of EEs in the mutant cells (Figure 1A, C-D). This was specific for EEs as we did not observe this clustering for LAMP1-positive lysosomes (Figure 1E, Sup. Figure 1A), while Rab11-positive recycling endosomes were present in the perinuclear region irrespective of the genotype (Figure 1F, Sup. Figure 1B), as previously reported for WT cells (Ren *et al*, 1998).

**Figure 1.**
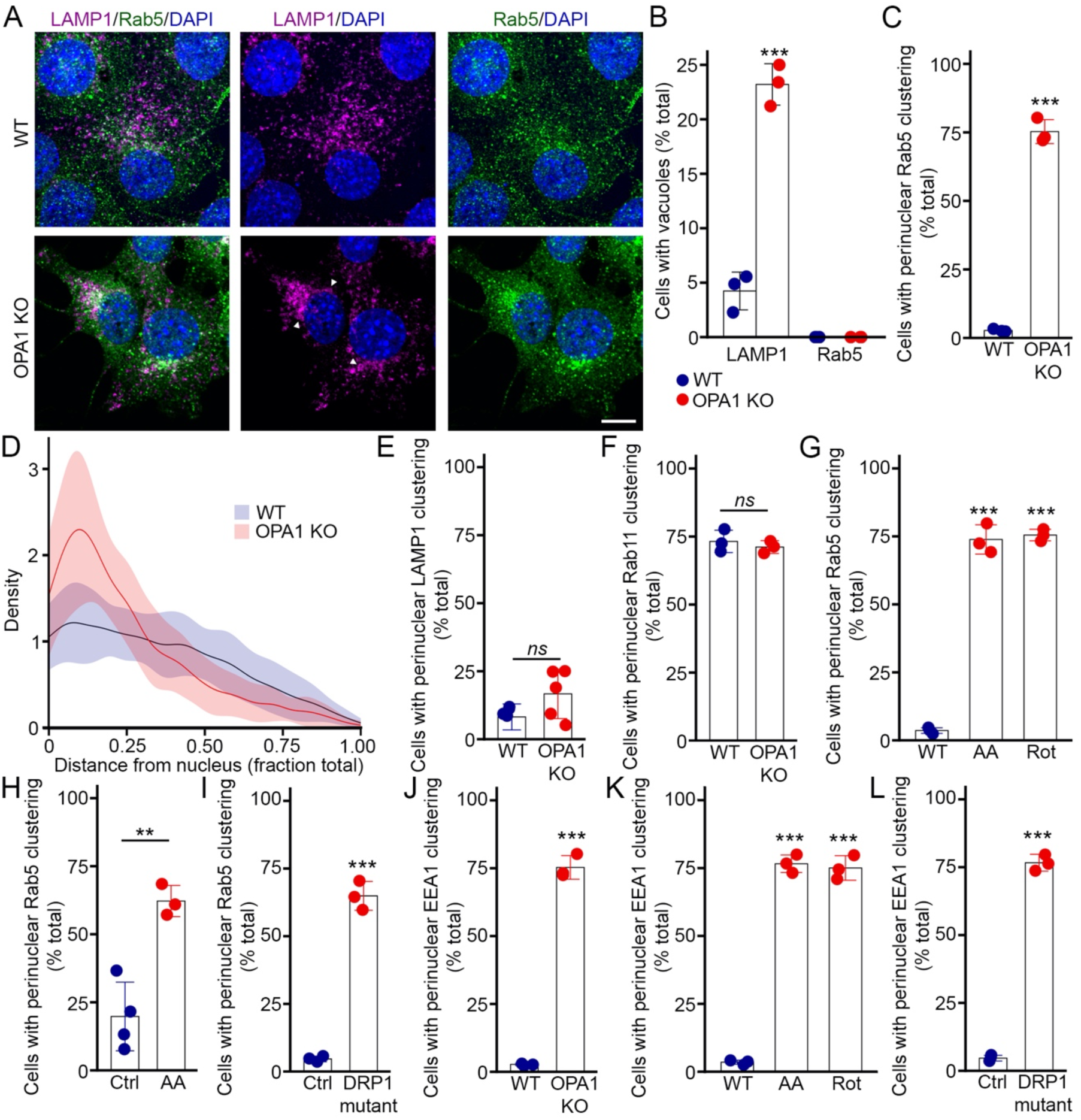
Mitochondrial dysfunction alters the distribution of early endosomes. (A) Representative confocal images of WT and OPA1 KO MEFs stained for the lysosomal marker LAMP1 (magenta) and the EE marker Rab5 (green), along with DAPI to mark nuclei (blue). Arrowheads denote enlarged vacuolated lysosomes. Scale bar 10 µm. (B) Quantification of WT and OPA1 KO cells with LAMP1-positive and Rab5-positive vacuoles from images in (A). Each point represents an independent experiment. Bars show the average ± SD for 3 experiments per condition. *** p<0.001, two-sided t-test. (C-F) Quantification of perinuclear clustering of the endosomal markers Rab5 (C), LAMP1 (E) and Rab11 (F) in OPA1 KO MEFs from confocal images as in (A). Each point represents an independent experiment. Bars show the average ± SD for 3 experiments per condition (5 for LAMP1). *** p<0.001, two-sided t-test. ns, not significant. The density of Rab5-positive vesicles relative to their localisation was also measured for Rab5 from the same images (D). The data shows the quantification of 30 cells per condition in 3 independent experiments ± SD. (G) Quantification of perinuclear clustering of Rab5 in WT MEFs treated with the Complex III inhibitor Antimycin A (AA) or the Complex I inhibitor Rotenone (Rot). Each point represents an independent experiment. Bars show the average ± SD for 3 experiments per condition. *** p<0.001, One-way ANOVA (H) Quantification of Rab5 perinuclear clustering in AA-treated HeLa cells. Each point represents an independent experiment. Bars show the average ± SD for 3 experiments per condition. ** p<0.01, two-sided t-test (I) Quantification of Rab5 perinuclear clustering in Control and DRP1 mutant primary fibroblasts. Each point represents an independent experiment. Bars show the average ± SD for 3 experiments per condition. *** p<0.001, two-sided t-test. (J-L) Quantification of perinuclear clustering of the EE marker EEA1 in OPA1 KO MEFs (J), WT MEFs treated with AA or Rotenone (K) and DRP1 mutant primary fibroblasts (L). Bars show the average ± SD for 3 experiments per condition. *** p<0.001, two-sided t-test, one-way ANOVA for (K).

To confirm that the alterations we observed in EEs are caused by the impairment of mitochondrial function, not some other defect associated with mutant fibroblasts, we chemically inhibited subunits of the electron transport chain (ETC) in WT MEFs. Similar to OPA1 KO MEFs, rotenone (Complex I inhibitor) or Antimycin A (AA, Complex III inhibitor) caused the perinuclear clustering of Rab5-positive EEs (Figure 1G). AA treatment also caused EE perinuclear clustering in HeLa cells (Figure 1H). Interestingly, a similar clustering of rab5-positive EEs was observed in primary human fibroblasts mutant for the mitochondrial fission protein DRP1 (Figure 1I). To confirm that the changes are the consequence of altered EEs, not simply a change in Rab5 distribution, we determined EE distribution using EEA1 as a second EE marker. Similar to Rab5-positive EEs, we found that EEA1-positive early endosomes clustered in the perinuclear region of OPA1 KO MEFs, WT MEFs treated with ETC inhibitors and DRP1 mutant fibroblasts (Figure 1J-L, Sup. Figure 1D), further indicating that EEs are altered in these cells.

### EE clustering in cells with mitochondrial dysfunction is associated with impaired EE cargo trafficking

The altered EE distribution in cells with mitochondrial dysfunction prompted us to address its consequences on EE function. EEs serve as sorting organelles to transfer cargo to late endosomes/lysosomes for degradation, or recycling endosomes for material returning to the cell surface (O’Sullivan & Lindsay, 2020). We first tracked the transfer of EE cargo towards lysosomes using fluorescently labelled dextran. Cells were first pulsed for 5 minutes with dextran, after which dextran uptake was evaluated by immunofluorescence and quantified by measuring the number of dextran-positive vesicles present in each cell. As cells with mitochondrial dysfunction showed similar uptake of dextran than control cells (Figure 2A), we then performed a chase experiment where the fate of the dextran is followed over time after the initial pulse. We investigated the trafficking of dextran in these endosomes by measuring the association of dextran with Rab5 and the lysosomal marker LAMP1 by confocal microscopy. As expected, dextran colocalised with Rab5-positive EEs immediately after the initial pulse, and gradually transferred to LAMP1-positive late endosomes/lysosomes over time in control cells (Figure 2B-C). We also confirmed this manual evaluation of colocalization by measuring Pearson correlation coefficients in the same images. While the overall low amount of colocalization between dextran and Rab5 caused low Pearson coefficient values, there was a clear increase in Pearson coefficient between dextran and LAMP1 over time, confirming our colocalization data (Sup. Figure 2A-B).

**Figure 2.**
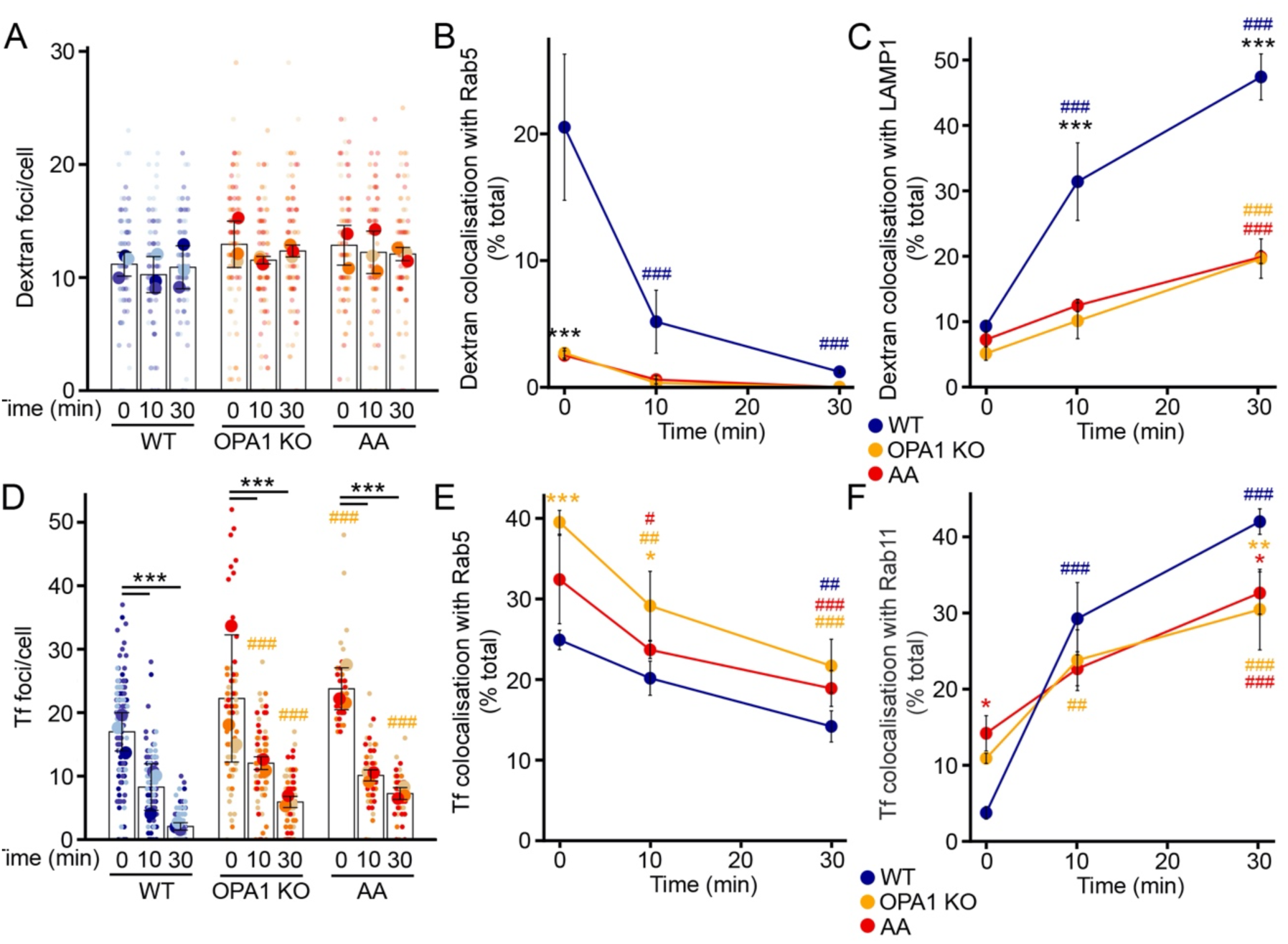
Mitochondrial dysfunction impairs proper EE cargo trafficking. (A-C) Dextran trafficking in OPA1 KO MEFs and AA-treated WT MEFs. Cells were pulsed with dextran for 5 minutes then chased for the indicated times and total dextran (A) and its colocalization with Rab5 (B) and LAMP1 (C) quantified by immunofluorescence. Each point represents an independent experiment, with small points in (A) representing individual cells. Bars show the average ± SD for 3 experiments per condition. *** p<0.001 vs WT, ### p<0.001 vs 0 min, two-way ANOVA. (D-F) Transferrin (Tf) trafficking in OPA1 KO MEFs and AA-treated WT MEFs. Cells were pulsed with Tf for 5 minutes then chased for the indicated times and total Tf (D) and its colocalization with Rab5 (E) and Rab11 (F) quantified by immunofluorescence. Each point represents an independent experiment, with small points in (D) representing individual cells. Bars show the average ± SD for 3 experiments per condition. *** p<0.001 vs WT, ### p<0.001 vs 0 min, two-way ANOVA.

Having validated the assay, we then measured dextran distribution in the endosomes of OPA1 KO MEFs. In contrast to WT cells, dextran colocalized with Rab5-positive EEs to a much lower extent in OPA1KO MEFs. Consistent with this, dextran transferred to LAMP1-positive lysosomes at a lower rate in OPA1 KO MEFs (Figure 2B-C, Sup. Figure 2A-B), demonstrating a defect in cargo delivery from the EEs to lysosomes in these cells. Similar results were obtained when WT MEFs were treated with AA (Figure 2B-C, Sup. Figure 2A-B).

We then determined the recycling function of EEs by tracking the delivery of cargo from EEs to recycling endosomes using fluorescently labelled transferrin (Tf). Tf binds to the Tf receptor at the cell surface, liberates its associated iron within EEs and is then recycled to the cell surface in recycling endosomes. As with dextran, an initial pulse of Tf led to the endocytosis of a similar amount of Tf in control, OPA1 KO and AA-treated MEFs, as judged by the similar number of Tf-positive vesicles present in these cells (Figure 2D). In control cells, the total number of transferrin spots then dropped, as expected because it is recycled to the cell surface (Figure 2D). Tf foci also decreased in OPA1 KO cells and WT cells treated with AA, although at a somewhat reduced rate (% left at 30 min: WT 12.5±4.05%; KO 28.9±8.24%, p=0.022; AA 30.5±1.62%, p=0.015). Interestingly, the colocalization with Rab5-positive EEs was somewhat increased in cells with mitochondrial dysfunction (Fig. 2E, Sup. Figure 2C). In addition, the colocalization between Tf and the recycling endosome marker Rab11 was somewhat increased at the beginning of the chase time but not at later time points (Fig. 2F, Sup. Figure 2D). Overall, these results suggest that endosomal recycling is partially affected in these cells. Thus, our results indicate that mitochondrial loss of function affect both endosomal recycling and delivery to lysosomes, although the latter is much more affected.

### EE clustering is driven by microtubule-dependent retrograde transport

Mitochondrial dysfunction induces perinuclear aggregation of EEs, resulting in altered EE spatial distribution that subsequently impacts cargo trafficking towards lysosomes. In order to elucidate the underlying cause, we initially determined the behaviour of EEs by live cell imaging. WT and OPA1 KO cells were transfected with RFP-Rab5 and imaged over time using a confocal microscope. We then analysed the movement of RFP-Rab5-positive EEs in these cells by measuring the radial (towards/away from the nucleus) and angular (side-to-side) components of vesicle speed (Figure 3A). This revealed that while the angular velocity of the EEs was not significantly affected by OPA1 deletion, their radial velocity was significantly increased (Figure 3B-C). Specifically, and consistent with the EE clustering we observed, mutant cells exhibited an augmented velocity of EEs directed towards the nucleus relative to control cells (Figure 3D). We then determined if these alterations were selective for EEs by repeating the experiment in cells in which lysosomes were tagged using GFP-LAMP1. In contrast to Rab5-positive vesicles, the radial velocity of LAMP1-positive vesicles was not altered in OPA1 KO MEFs (Figure 3B, D), although a subset of cells showed increased angular velocity (Figure 3C). Altogether, these results indicate that EE movements are increased in OPA1 KO cells.

**Figure 3.**
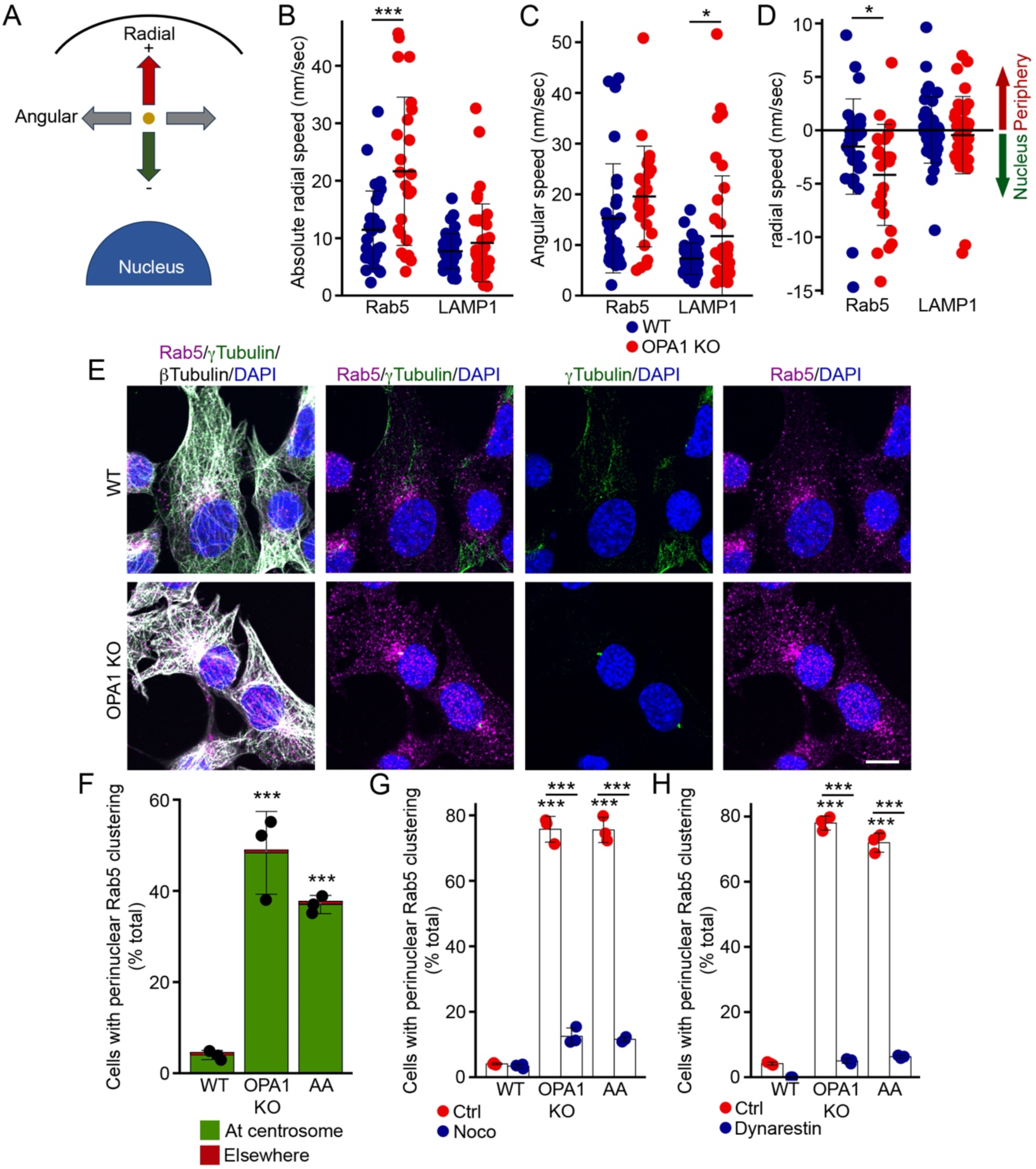
Microtubule drive EE perinuclear clustering. (A-D) Quantification of EE and lysosome velocity in OPA1 KO MEFs transfected with either Rab5-RFP (EE) or LAMP1-GFP (lysosome). (A) schematic representation of radial and angular velocity. (B) Absolute radial speed (independent of direction), (D) Angular speed, (E) Radial speed according to its direction (negative towards the nucleus, positive towards the cell membrane). Each point represents an individual endosome, with endosomes analysed in a minimum of 4 cells per condition in at least 3 independent experiments. Bars show the average ± SD. *** p<0.001, * p<0.05, two-sided t-test. (E-F) Rab5-positive EEs cluster at centrosomes. (D) Representative confocal images of WT and OPA1 KO MEFs stained for the EE marker Rab5 (green), along with the microtubule marker ß-tubulin and the centrosome marker γ-tubulin. Scale bar 10 µm. (E) Quantification of images in (D). Each point represents an independent experiment. Bars show the average ± SD for 3 experiments per condition. *** p<0.001, one-way ANOVA. (G-H) Inhibition of microtubule-dependent transport rescues Rab5-positive EE distribution in OPA1 KO MEFs. WT and OPA1 KO MEFs were treated with the microtubule inhibitor nocodazole (noco, G) or the the Dynein inhibitor Dynarestin (H) in the absence or the presence of AA as indicated. Each point represents an independent experiment. Bars show the average ± SD for 3 experiments per condition. *** p<0.001, one-way ANOVA.

As EE transport occurs along microtubules, we next investigated the relationship between EE clustering and microtubule transport in cells with mitochondrial dysfunction. For this, we first stained WT and OPA1 KO cells for Rab5, the microtubule protein ß-tubulin and the centrosome marker γ-tubulin. In control cells, where EEs were dispersed throughout the cell, no association between EEs and centrosomes was observed (Figure 3E-F). Conversely, in mutant cells characterized by the aggregation of early endosomes in the perinuclear region, we identified the minus end of microtubules and centrosomes associating with the aggregated early endosomes (Figure 3E-F). This triple association substantiates the involvement of microtubules and centrosomes in the aggregation of early endosomes.

To investigate the causal relationship between microtubules and the altered distribution of early endosomes, we employed the microtubule depolymerizing agent nocodazole as an experimental tool. A short pulse of nocodazole (15 min) had no impact on the distribution of Rab5-positive EEs in control cells (Figure 3G). In contrast, nocodazole treatment dispersed aggregated EEs from the perinuclear region in mutant cells (Figure 3G). A similar redistribution of Rab5 upon nocodazole treatment was observed in WT cells treated with AA (Figure 3G), consistent with microtubule-dependent transport being required for perinuclear EE clustering. The motor protein Dynein plays a key role in the retrograde transport of EE (Schuster *et al*, 2011). Thus, we then determined the effect of the Dynein inhibitor Dynarestin on Rab5 clustering in cells with mitochondrial dysfunction. Similar to the disruption of microtubules with nocodazole, Dynarestin treatment caused a redistribution of Rab5-positive EEs in OPA1 KO cells and in AA-treated WT cells (Figure 3H). Altogether, these results indicate that microtubule-based transport promotes perinuclear EE aggregation in cells exhibiting mitochondrial dysfunction.

### Mitochondrial dysfunction leads to centrosome alterations

To determine the reason why microtubule-based transport specifically altered EE localisation in cells with mitochondrial dysfunction, we first assessed potential alterations in microtubule organisation. Most interphase cells contain one centrosome from where microtubules are organised, while dividing cells have duplicated their centrosome in prevision of chromosome segregation and cytokinesis (Prigent & Uzbekov, 2022). Consistent with this, almost all WT MEFs contained either one (57%) or two (41%) centrosomes (Figure 4A-B). In contrast, OPA1 KO cells and AA-treated WT cells displayed a decrease in cells with one centrosome, and a significant proportion of cells with more than two centrosomes (Figure 4A-B). In addition, while the two centrosomes were apart from each other in WT cells, as would be expected from cells preparing to divide, the distance between the two centrosomes decreased as the number of centrosomes increased in OPA1 KO and AA-treated cells (Figure 4C), suggesting that duplicated centrosomes fail to segregate in these cells.

**Figure 4.**
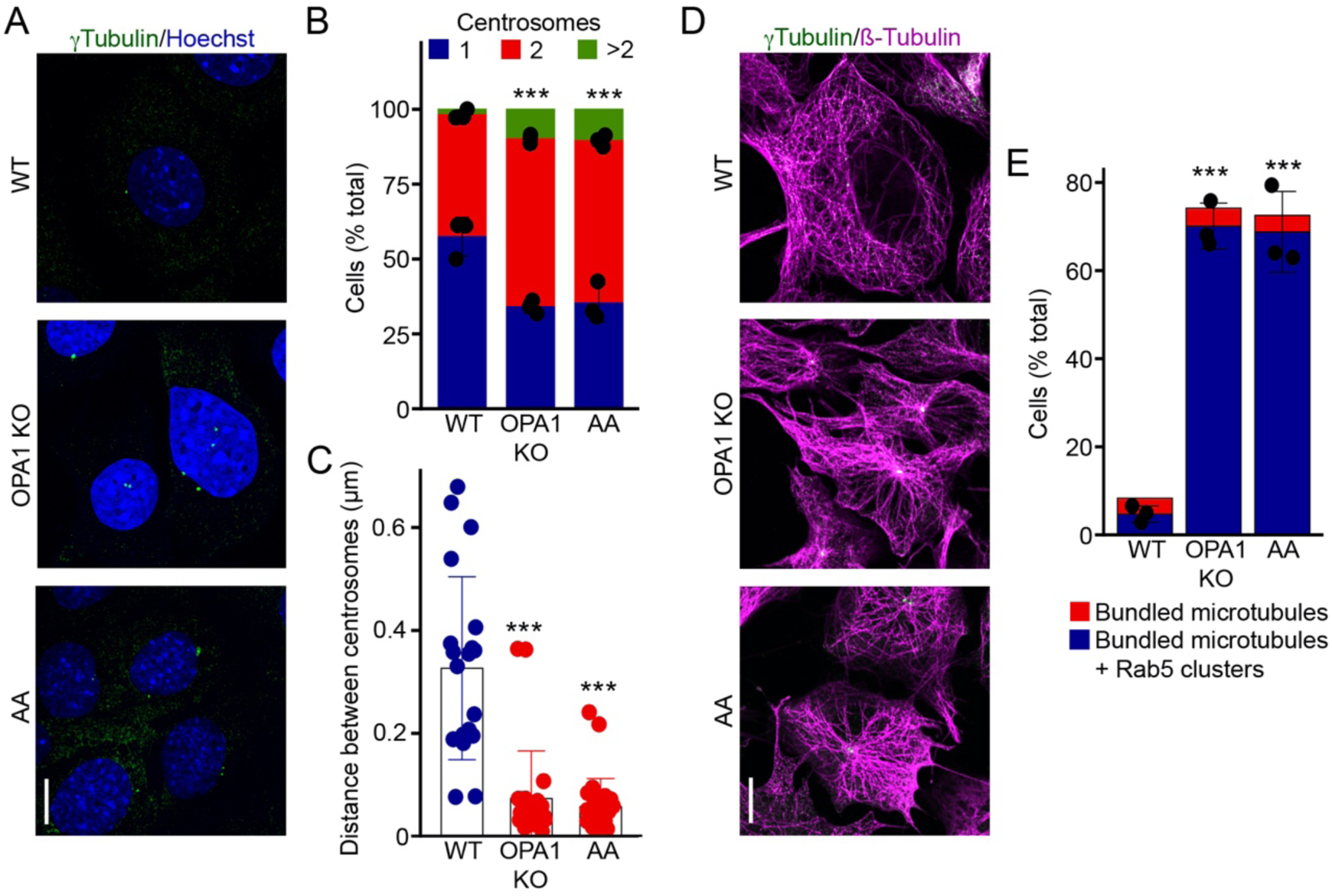
Mitochondrial dysfunction leads to aberrant centrosome duplication. (A) Representative confocal images of WT (control and AA-treated) and OPA1 KO MEFs stained for the centrosome marker γ-tubulin (green) along with DAPI (blue) to mark nuclei. Scale bar 10 µm. (B) Quantification of the number of centrosomes per cell from images in (A). Each point represents an independent experiment. Bars show the average ± SD for 3 experiments per condition. *** p<0.001, two-way ANOVA. (C) Quantification of the distance between centrosomes from images in (A). Each point represents an independent experiment. Bars show the average ± SD for 3 experiments per condition. *** p<0.001, one-way ANOVA. (D) Representative confocal images of WT (control and AA-treated) and OPA1 KO MEFs stained for the centrosome marker γ-tubulin (green) and the microtubule marker ß-tubulin (magenta). Scale bar 10 µm. (E)

In osteoclasts, centrosome clustering facilitates microtubule bundling, thereby enhancing microtubule-based transport (Philip *et al*, 2022). A similar bundling was present in OPA1 KO MEFs and AA-treated WT MEFs, as seen by the presence of much thicker microtubule fibres in cells with closely apposed duplicated centrosomes compared to control cells (Figure 4D-E). Importantly, the presence of these dense microtubules correlated with EE perinuclear clustering (Figure 3E, 4E). Collectively, our results suggest that alterations in centrosomes lead to changes in the organization of microtubules, promoting rapid movement and aggregation of early endosomes driven by dynein in the perinuclear region.

### Oxidative stress promotes centrosome duplication and EE clustering

Mitochondrial dysfunction generally leads to an increase in reactive oxygen species (ROS)(Kausar *et al*, 2018) and we previously showed that this was associated with the impaired lysosomal structure and function found in these cell (Demers-Lamarche *et al*., 2016). Importantly, oxidative stress also causes aberrant centrosome duplication, leading to the presence of supernumerary centrosomes and aberrant cell division (Chae *et al*, 2005). Consistent with this, addition of glucose oxidase (GO) to cell media, to generate a controlled amount of ROS in WT cells (Demers-Lamarche *et al*., 2016), caused a significant increase in the number of cells with more than two centrosomes, as well as an increase in the number of cells with bundled microtubules (Figure 5A-C). Exposure of WT cells to GO also caused the perinuclear clustering of Rab5-positive EEs (Figure 5D), similar to what we observed in cells with mitochondrial dysfunction.

**Figure 5.**
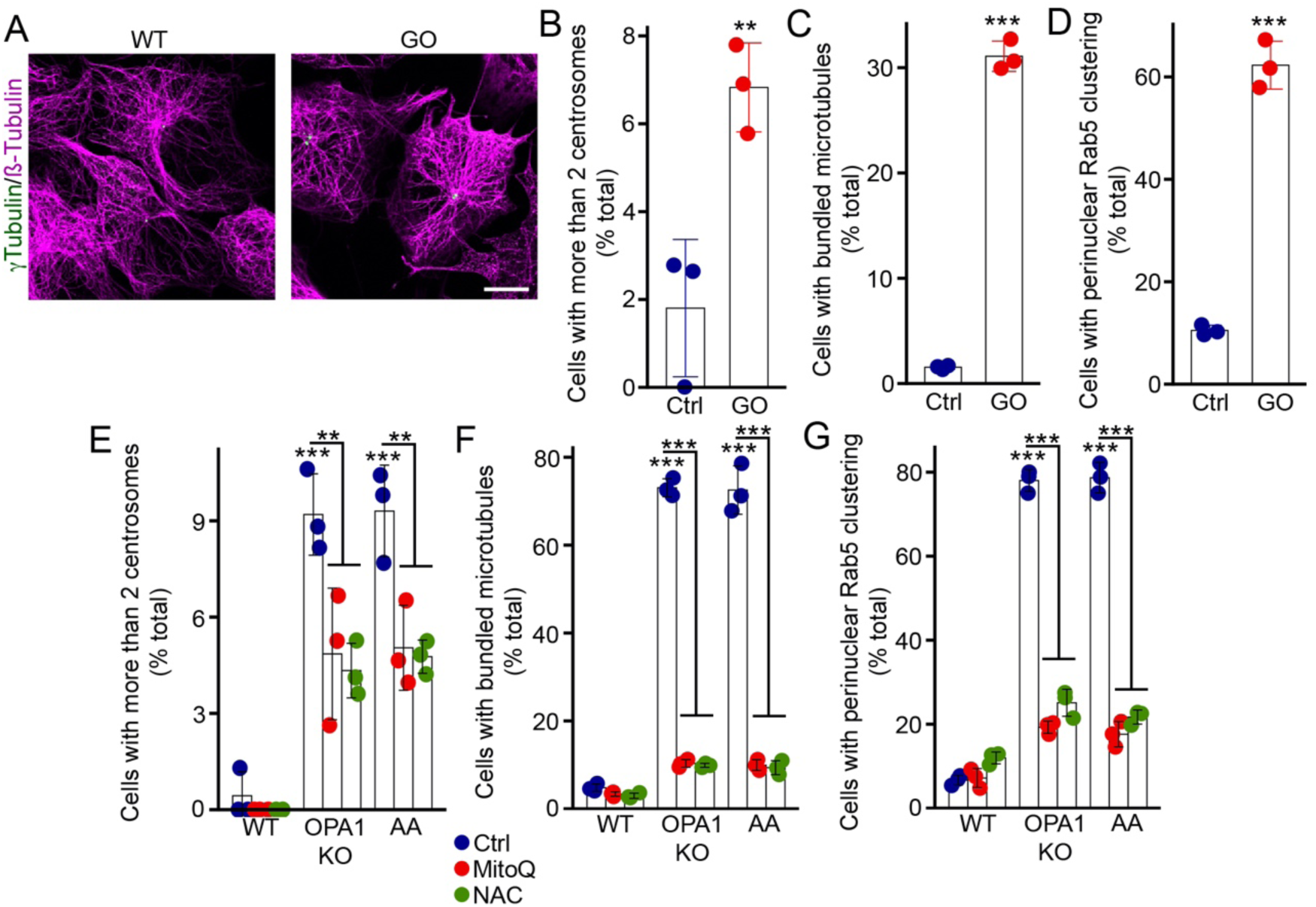
Oxidative stress promotes centrosome alterations leading to EE perinuclear clustering. (A-D) Glucose oxidase (GO) promotes centrosome alterations and EE clustering. (A) Representative confocal images of control and GO-treated WT MEFs stained for the microtubule marker ß-tubulin and the centrosome marker γ-tubulin. Scale bar 10 µm. The images were quantified for the presence of extra centrosomes (more than 2)(B), bundled microtubules (C) and Rab5-positive EEs perinuclear clustering (D). Each point represents an independent experiment. Bars show the average ± SD for 3 experiments per condition. *** p<0.001, ** p<0.01, two-sided t-test. (E-G) Antioxidants rescue both centrosomes defects and EE aggregation. WT and OPA1 KO MEFs were treated as indicated and the presence of extra centrosomes (more than 2)(E), bundled microtubules (F) and Rab5-positive EEs perinuclear clustering (G) were quantified from confocal images as in (A-D). Bars show the average ± SD for 3 experiments per condition. *** p<0.001, ** p<0.01, two-way ANOVA.

To further demonstrate that ROS cause the centrosome alterations leading to EE clustering, we quenched ROS in cells with mitochondrial dysfunction using the antioxidants MitoQ and N-Acetyl-cystein (NAC). Consistent with our GO data, antioxidants decreased the number of OPA1 KO and AA-treated cells that contained more than 2 centrosomes (Figure 5E). Similarly, we observed a rescue of the dense microtubule phenotype near the minus end of the microtubule in antioxidant-treated cells (Figure 5F). This rescue of microtubule structure caused the dispersion of EEs from the perinuclear region towards the entire cell (Figure 5G), supporting a key role for ROS in this process. Collectively, our data demonstrates that oxidative stress-induced changes in microtubule organization and centrosomes contribute to the altered EE distribution.

### Oxidative stress causes a functional loss of cargo trafficking

As ROS promote EE clustering, we then investigated the impact of oxidative stress on the trafficking capacity of EEs. We first exposed control cells to GO to induce an increase in ROS production. Consistent with our models of mitochondrial dysfunction, GO did not decrease the overall uptake of dextran (Figure 6A). However, GO reduced the colocalization between dextran and Rab5-positive EEs and reduced trafficking towards LAMP1-positive lysosomes (Figure 6B-E), similar to cells with mitochondrial dysfunction (Figure 2C-D). We then exposed OPA1 KO and AA-treated cells to the antioxidant MitoQ, which significantly reduced the microtubule alterations in these cells (Figure 5E-F). Consistent with oxidative stress playing a key role in the EE alterations present in cells with mitochondrial dysfunction, MitoQ treatment had no impact on total dextran uptake (Figure 6A), but rescued dextran transport to Rab5-positive EEs and LAMP1-positive lysosomes (Figure 6D-G). Altogether, our results demonstrate that ROS alter EE distribution and subsequent cargo trafficking to lysosomes.

**Figure 6.**
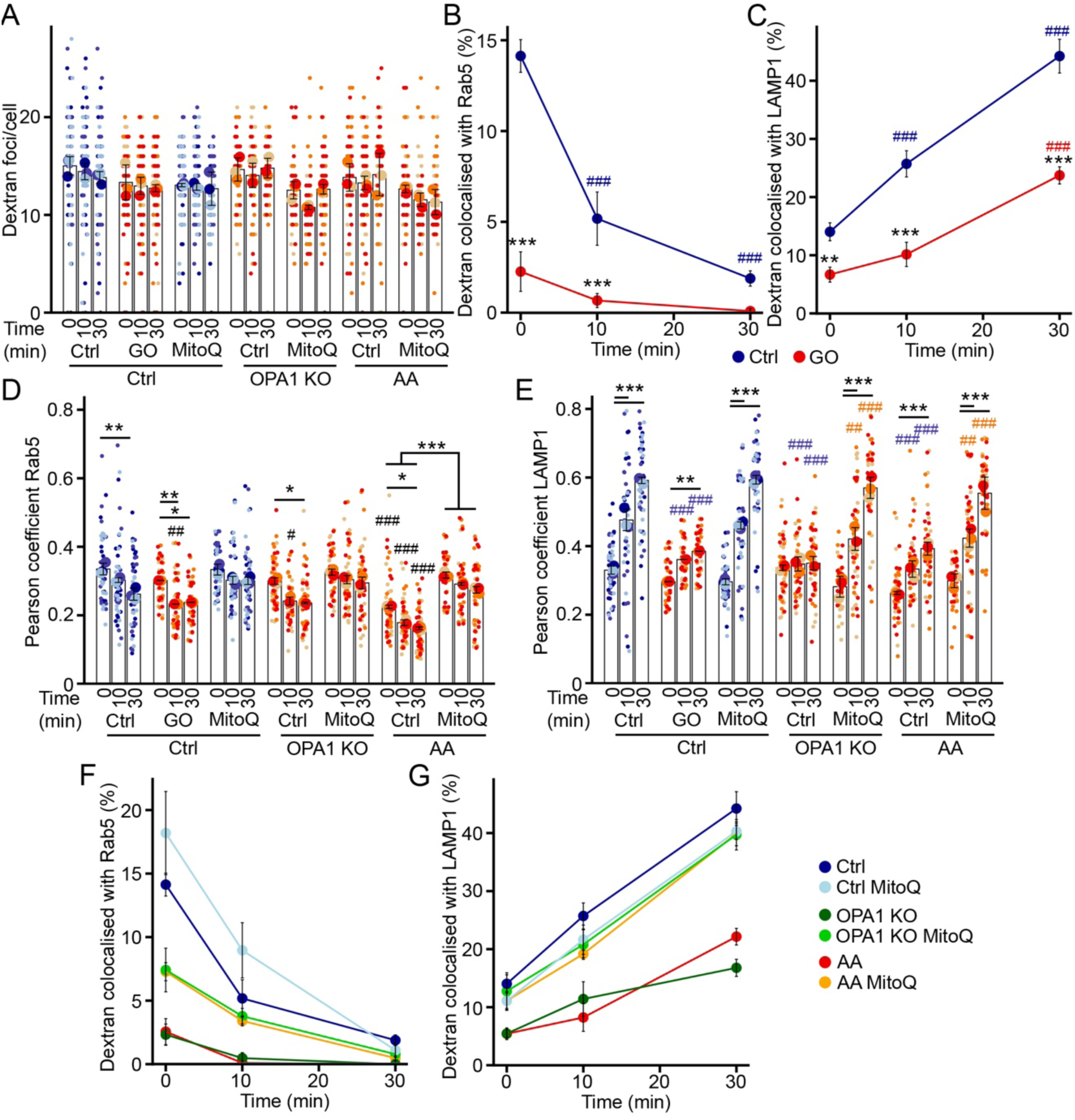
Antioxidants rescue EE function in cells with mitochondrial dysfunction. (A-C) Dextran trafficking in WT MEFs treated with glucose oxidase (GO). Cells were pulsed with dextran for 5 minutes then chased for the indicated times and total dextran (A) and its colocalization with Rab5 (B) and LAMP1 (C) quantified by immunofluorescence. Each point represents an independent experiment, with small points in (A) representing individual cells. Bars show the average ± SD for 3 experiments per condition. *** p<0.001 vs WT, ### p<0.001 vs 0 min, two-way ANOVA. (D-F) Antioxidants rescue dextran trafficking in OPA1 KO MEFs and AA-treated WT MEFs. Cells were pulsed with dextran for 5 minutes then chased for the indicated times and total dextran (D) and its colocalization with Rab5 (E) and LAMP1 (F) quantified by immunofluorescence. Each point represents an independent experiment, with small points in (D) representing individual cells. Bars show the average ± SD for 3 experiments per condition. *** p<0.001 vs WT, ### p<0.001 vs 0 min, two-way ANOVA.

## Discussion

Mitochondria interacts with various organelles to maintain cellular homeostasis. While functional interactions with the endoplasmic reticulum and lysosomes are well established (Zhang *et al*, 2023), the influence of mitochondria on other components of the endosomal system is less understood. Here, we addressed this question using genetic and chemical models of mitochondrial dysfunction. Our findings revealed significant alterations in the distribution and function of EEs in cells with impaired mitochondrial activity. Notably, these changes are caused by ROS-dependent alterations in centrosomes and microtubules affecting EE transport. The impaired distribution of EEs observed following mitochondrial dysfunction affected their ability to efficiently deliver cargo to lysosomes.

Several reports have shown that mitochondrial activity is required to maintain functional lysosomes, while impaired lysosomes cause the accumulation of damaged mitochondria as a result of blocked mitophagy (Deus *et al*, 2020; Stepien *et al*, 2020). Thus, a complex interplay exists between the two organelles to control energy production, metabolite availability and quality control processes. Interestingly, we previously reported that the inhibition of lysosomal activity in cells with dysfunctional mitochondria depends on ROS production (Demers-Lamarche *et al*., 2016). Here, we show that mitochondrial dysfunction also leads to ROS-dependent early endosome aggregation, suggesting that the consequences of mitochondrial dysfunction are not limited to lysosomes but also affect early endosomes. This impairs cargo trafficking from early endosomes to lysosomes, which could contribute to the lysosomal defects present in these cells.

In physiological conditions, mitochondrial ROS play essential roles in cellular signalling (Mailloux, 2020). Nevertheless, an imbalance in ROS production and removal can lead to oxidative stress (Zorov *et al*, 2014). Alterations in mitochondrial structure and function caused by mutations or inhibition of the electron transport chain, as well as the accumulation of damaged mitochondria following the inhibition of mitophagy, can dramatically increase the production of mitochondrial ROS (Zong *et al*, 2024). These ROS can then cause cellular damage and contribute to the development of neurodegenerative, metabolic, cardiovascular, and inflammatory diseases (Chae *et al*., 2005; Kumar *et al*, 2012; Madamanchi & Runge, 2007; Silwal *et al*, 2020; Tirichen *et al*, 2021).

In the context of organelle trafficking and function, one of the important targets of ROS is the cytoskeleton. For example, exposure of cells to high concentrations of H_2_O_2_ can cause microtubule depolymerization and loss of architectural stability (Caporizzo & Prosser, 2022; Goldblum *et al*, 2021). However, the lower levels of endogenous ROS produced in our models of mitochondrial dysfunction or GO treatment (Demers-Lamarche *et al*., 2016), did not cause microtubule depolymerization. Rather, this oxidative stress was associated with the presence of supernumerary centrosomes, consistent with a previous report indicating that ROS promote centrosomes amplification (Chae *et al*., 2005). These centrosomal clusters increase microtubule nucleation and alter intracellular trafficking under physiological conditions in osteoclasts (Philip *et al*., 2022). We similarly found that the ROS-dependent centrosome amplification present in cells with mitochondrial dysfunction altered EE trafficking, causing their accumulation around these centrosomes. As for the absence of detectable redistribution of lysosomes in cells with defective mitochondria, it is likely the consequence of the more complex trafficking of these organelles, both anterograde and retrograde, in contrast to the dominant retrograde transport of EEs from the plasma membrane.

Pathological conditions due to mitochondrial defects were originally explained by defects in ATP production. However, more recent work has highlighted the multifaceted aspects of mitochondria-related diseases, including several metabolic alterations not directly related to ATP production, ROS production and alterations in organelle contact sites (Deus *et al*., 2020; Ilaria *et al*, 2024; Tan & Finkel, 2020). This complex interplay between metabolism and organelle distribution and activity likely plays a major role in the etiology of mitochondria-related diseases. For example, neurodegenerative diseases are associated with defects in both mitochondria and the endolysosomal compartment, with oxidative stress playing an important role (Deus *et al*., 2020; Patten *et al*, 2010). In this context, it is noteworthy that we identified microtubule transport as an important target of ROS that impact EE function and could explain some of the features of these diseases.

Overall, we propose that the elevated ROS generated by damaged mitochondria cause aberrant centrosome duplication, leading to microtubule alterations that alter EE trafficking and impair their ability to transfer their cargo to lysosomes. Our model is supported by the fact that EE and microtubule defects are rescued by antioxidants and can be recapitulated in healthy cells by adding a controlled source of ROS (GO). Our study highlights that mitochondrial dysfunction not only impacts lysosomes but also influences the function and distribution of EEs, likely contributing to the neuronal impairment caused by mitochondrial dysfunction in neurodegenerative diseases.

## Methods

Cell culture reagents were bought from Wisent. Other chemicals were purchased from Sigma-Aldrich, except where indicated.

### Cell culture

WT and OPA1 KO MEFs (gift from Dr. Luca Scorrano, University of Padua), Control and DRP1 mutant primary human fibroblasts (Ilamathi *et al*, 2023), and HeLa were cultured in Dulbecco’s modified Eagle’s medium (DMEM) supplemented with 10% fetal bovine serum. Cells were maintained in an incubator with 5% CO2 until they reached 70-80% confluency before commencing the experiments. Cells were treated as follow: mitochondrial inhibitors antimycin A (50 μM) or rotenone (5 μM) for 4 hours; antioxidants N-acetylcysteine (NAC) (10 mM) or MitoQ (100 μM, Focus Biomolecules # 10-1363) for 4 hours; induction of oxidative stress with 50 milliunits/ml of glucose oxidase for 1 hour; microtubule depolymerisation with nocodazole (5 μM) for 15 minutes; inhibition of dynein with dynarrestin (2 μM) for 30 minutes.

### Dextran and transferrin uptake experiments

MEFs were grown on coverslips until they reached 70% confluency. Cells were then exposed to 0.5 mg/ml of Dextran, Tetramethylrhodamine, 10,000 MW, Lysine Fixable (fluoro-Ruby) (Thermofischer # D1868) for 5 minutes. The media containing dextran was then removed, and fresh media was added. The cells were then chased for 0, 10, and 30 minutes before being fixed with 4% paraformaldehyde (PFA). For the Tf experiments, the cells were kept in serum-free media on ice for 30 minutes, then incubated with 0.25 mg/ml of Transferrin from Human Serum (Tetramethylrhodamine Conjugate, Thermofisher #T2872) for 5 minutes. Afterward, the media containing transferrin was replaced with fresh media, and the cells were chased for 0, 10, and 30 minutes before being fixed 10 minutes with 4% PFA at 25° C. To investigate the endosomal/lysosomal trafficking function under oxidative stress conditions, the cells were first treated with glucose oxidase or MitoQ as above. The media was then changed, and cells were exposed to dextran as above.

### Immunofluorescence and cell imaging

Cells were grown on glass coverslips for 24 hours before the experiments, then treated as indicated and fixed 10 min with 4% PFA at 25° C. Cells were permeabilized with 0.2% Triton X-100 in PBS and blocked with 1% BSA / 0.1% Triton X-100 in PBS. Cells were then incubated with primary antibodies followed by fluorescent tagged secondary antibodies (Jackson Immunoresearch, 1:500) and DAPI (Invitrogen, Thermo Fisher, D1306, 1:100). The following primary antibodies were used: mouse Anti-β-Tubulin (Sigma-Aldrich # T5293), rat Anti-β-Tubulin (Abcam #6161), mouse anti-gamma tubulin (Sigma-Aldrich #T5326), rat anti-LAMP1 (SCBT #19992), rabbit anti-Rab5 (Cell signalling Technologies #3547S, 1:200), rabbit anti-EEA1 (Cell signalling Technologies #2411, 1:200), and rabbit anti-Rab11 (Cell Signaling Technologies #3539S, 1:200). Imaging was performed using a Leica TSC SP8 confocal microscope fitted with a 63×/1.40 oil objective.

For live cell imaging, WT and OPA1 KO MEFs were transfected with mRFP-Rab5 (Addgene plasmid # 14437) or LAMP1-GFP (Addgene plasmid # 16290) using Metafecten Pro (Biontex). 24h later the cells were plated on glass bottom dishes in complete medium and grown for another 24h. The plates were then mounted onto Leica TSC SP8 confocal microscope fitted with a 63×/1.40 oil objective. Time-lapse images were acquired at a speed of (0.05-0.125 frames/s) for 10 minutes.

### Image processing and analysis

All image manipulation and analysis were done in Image J/Fiji. The images shown are from single focal planes unless stated otherwise. For density distributions, the images were first segmented in ImageJ using Filter/Median (1.0), then thresholding and adjusting the resulting image using Binary/Erode. The position of each foci was then identified using the Analyse particle function with Size set to 0.75 µm-Infinity. The position of the nucleus and the plasma membrane was also identified. The data was then fed to a custom R script to calculate the relative distance to the nucleus and the distribution calculated using the Density function. Radial and Angular speeds were similarly calculated using a custom R script.

### Data analysis and statistics

Data analysis and statistical procedures were conducted using R. Quantification of immunofluorescence data was performed, and representative images from a minimum of three independent experiments were presented (specific sample sizes are indicated in the respective quantification figures). Data is expressed as the mean ± standard deviation (SD) as indicated in the figure legends. To assess statistical significance, Student’s t-test was utilized for comparisons between two groups, while one-way ANOVA with a Tukey post hoc test was employed for multiple comparisons.

## Acknowledgements

This work was supported by grants from the Natural Sciences and Engineering Research Council of Canada and the Fondation UQTR. K.T. was a recipient of a Queen Elizabeth II Diamond Jubilee scholarship and a Fonds du Québec-Santé scholarship. L.C. was a recipient of a Fonds du Québec-Santé scholarships.

---

**Supplementary Figure 1.**
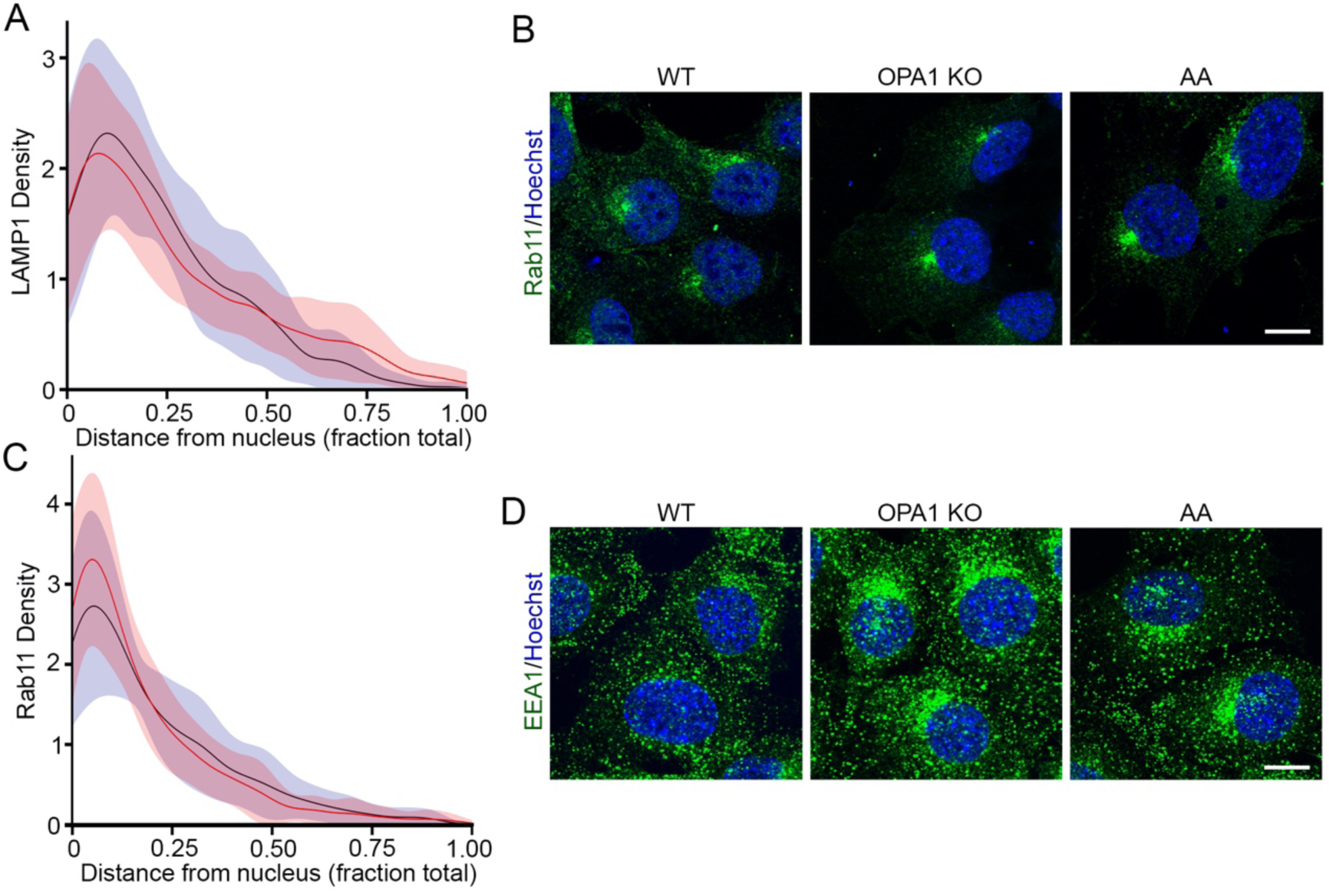
Distribution of endosomal markers in cells with mitochondrial dysfunction. (A) Density of LAMP1-positive vesicles relative to their localisation as measured from confocal images in Figure 1A. The data shows the quantification of 30 cells per condition in 3 independent experiments ± SD. (B) Representative confocal images of WT, OPA1 KO and AA-treated WT MEFs stained for the recycling endosome marker Rab11 (green), along with DAPI to mark nuclei (blue). Scale bar 10 µm. (C) Density of Rab11-positive vesicles relative to their localisation as measured from confocal images in (B). The data shows the quantification of 30 cells per condition in 3 independent experiments ± SD. (D) Representative confocal images of WT, OPA1 KO and AA-treated WT MEFs stained for the EE marker EEA1 (green), along with DAPI to mark nuclei (blue). Scale bar 10 µm.

**Supplementary Figure 2.**
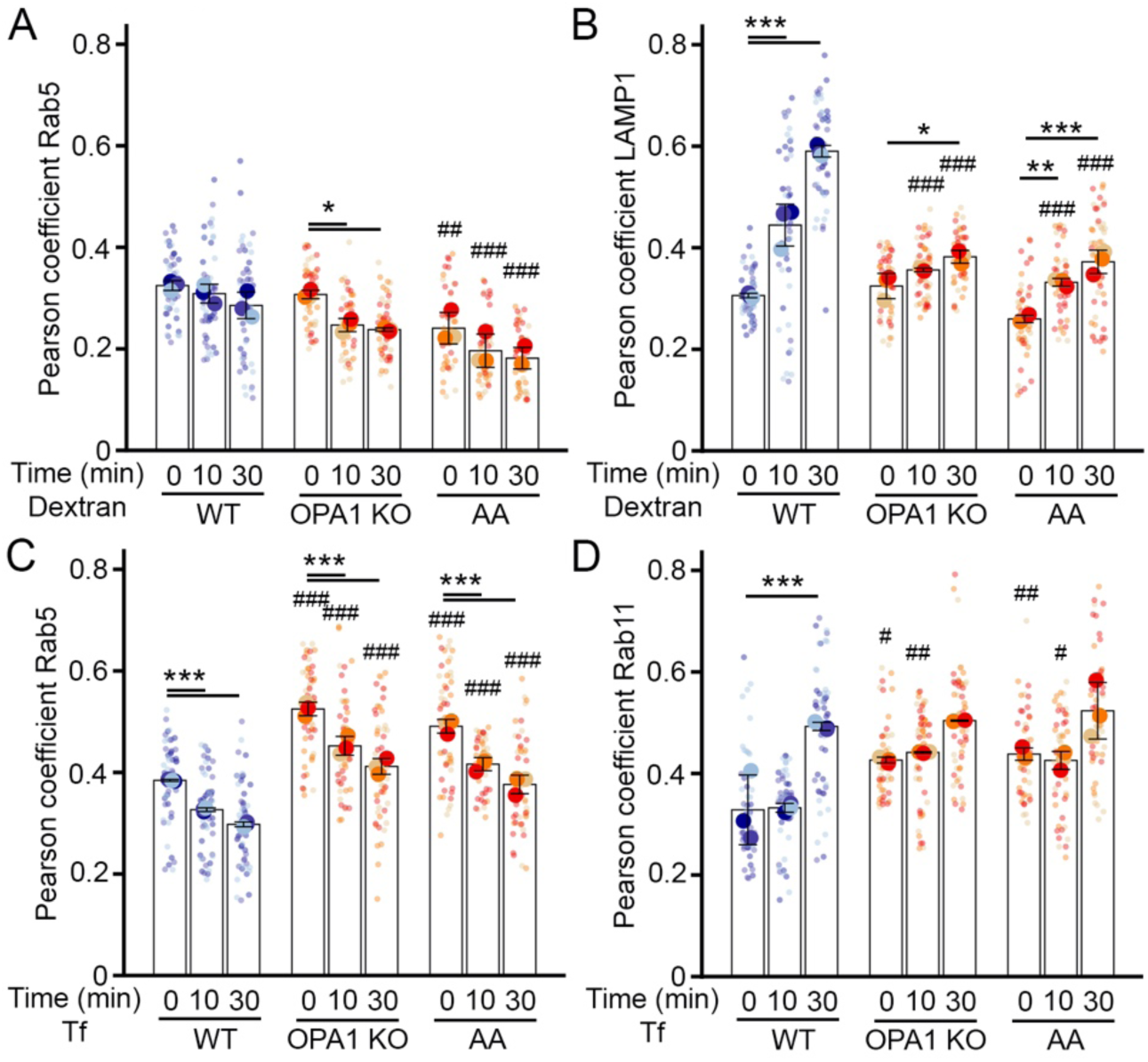
Loss of colocalisation between dextran and lysosomes in cells with mitochondrial dysfunction. (A-B) Dextran trafficking in OPA1 KO MEFs and AA-treated WT MEFs. Cells were pulsed with dextran for 5 minutes then chased for the indicated times and Pearson coefficients were calculated betewwn Dextran and Rab5 (A) or LAMP1 (B) from the same images as in Figure 2. Each point represents an independent experiment, with small points representing individual cells. Bars show the average ± SD for 3 experiments per condition. *** p<0.001, ** p<0.01, * p<0.05 vs WT, ### p<0.001, ## p<0.01 vs 0 min, two-way ANOVA. (C-D) Transferrin (Tf) trafficking in OPA1 KO MEFs and AA-treated WT MEFs. Cells were pulsed with Tf for 5 minutes then chased for the indicated times and Pearson coefficients were calculated betewwn Tf and Rab5 (C) or LAMP1 (D) from the same images as in Figure 2. Each point represents an independent experiment, with small points representing individual cells. Bars show the average ± SD for 3 experiments per condition. *** p<0.001 vs WT, ### p<0.001, ## p<0.01, # p<0.05 vs 0 min, two-way ANOVA.

